# SARS-CoV-2 outbreak in lions, tigers and hyenas at Denver Zoo

**DOI:** 10.1101/2024.10.14.617443

**Authors:** Emily N Gallichotte, Laura Bashor, Katelyn Erbeck, Lara Croft, Katelyn Stache, Jessica Long, Sue VandeWoude, James G Johnson, Kristy L. Pabilonia, Gregory D Ebel

## Abstract

In late 2019, SARS-CoV-2 spilled-over from an animal host into humans, where it efficiently spread, resulting in the COVID-19 pandemic. Through both natural and experimental infections, we learned that many animal species are susceptible to SARS-CoV-2. Importantly, animals in close proximity to humans, including companion, farmed, and those at zoos and aquariums, became infected, and many studies demonstrated transmission to/from humans in these settings. In this study, we first review the literature of SARS-CoV-2 infections in tigers and lions, and compare species, sex, age, virus and antibody detection assay, and types, frequency and length of clinical signs, demonstrating broad heterogeneity amongst infections. We then describe a SARS-CoV-2 outbreak in lions, tigers and hyenas at Denver Zoo in late 2021. Animals were tested for viral RNA (vRNA) for four months. Lions had significantly more viral RNA in nasal swabs than both tigers and hyenas, and many individual lions experienced viral recrudescence after weeks of undetectable vRNA. Infectious virus was correlated with high levels of vRNA and was more likely to be detected earlier during infection. Four months post-infection, all tested animals generated robust neutralizing antibody titers. Animals were infected with Delta lineage AY.20 identical to a variant circulating at less than 1% in Colorado humans at that time, suggesting a single spillover event from an infected human spread within and between species housed at the zoo. Better understanding of epidemiology and susceptibility of SARS-CoV-2 infections in animals is critical to limit the current and future spread and protect animal and human health.

**Importance:** Surveillance and experimental testing have shown many animal species, including companion, wildlife, and conservatory, are susceptible to SARS-CoV-2. Early in the COVID-19 pandemic, big cats at zoological institutions were among the first documented cases of naturally infected animals; however, challenges in the ability to collect longitudinal samples in zoo animals have limited our understanding of SARS-CoV-2 kinetics and clearance in these settings. We measured SARS-CoV-2 infections over three months in lions, tigers and hyenas at Denver Zoo, and detected viral RNA, infectious virus, neutralizing antibodies, and recrudescence after initial clearance. We found lions had longer and higher levels of virus compared to the other species. All animals were infected by a rare viral lineage circulating in the human population, suggesting a single spillover followed by interspecies transmission. These data are important in better understanding natural SARS-CoV-2 spillover, spread and infection kinetics within multiple species of zoo animals.

## Introduction

SARS-CoV-2 emerged in 2019 and rapidly spread across the globe resulting in a pandemic, causing more than 7 million deaths, strain on healthcare systems, and negative social, economic and cultural effects. SARS-CoV-2 has infected a wide range of animal species via natural and experimental infection [1]. Despite its animal origins, SARS-CoV-2 spreads efficiently among humans. Multiple instances of spillback from humans back into animals have been observed, including white-tailed deer, companion animals (cats and dogs), and mink [2]. Studies have shown that SARS-CoV-2 rapidly evolves following infection of new species, suggesting adaptation to new hosts [3-5]. Cross-species infection resulting in emergence of novel variants is one hypothesis for the genesis of variants of concern with the potential for increased transmission, disease, and immune evasion in humans or other species. [6].

In April 2020, the first felid infections of SARS-CoV-2 were reported in tigers and lions at the Bronx Zoo, New York [7-9]. Shortly thereafter, domestic cats were reported as the first companion animals with naturally occurring SARS-CoV-2 infections [10]. Since then, domestic and non-domestic felid infections have become increasingly common and have been reported across the globe [11]. Across all cats (domestic and non-domestic), there are a range of clinical signs and disease presentations observed [11, 12]. In most cases of naturally infected domestic cats, it was determined that the cats acquired the virus from cohabitating infected humans [13]. Conversely, in many documented cases of non-domestic felid infections, primarily in lions and tigers in zoological settings, the source of infection is unknown, with potential sources including animal care staff, zoo visitors, or other infected animals [13]. Several commercial vaccines have been developed for SARS-CoV-2 for use in animals (primarily in zoos) since early 2021 [14]. Despite infection prevention practices and policies implemented at zoos and other settings (e.g., mink farms), including personal protective equipment worn by staff, limiting visitors, vaccination of animals, etc., SARS-CoV-2 infections of animals in these settings have continued to be reported [15].

Many studies describing SARS-CoV-2 infections in non-domestic felids are limited by samples from single time points, and/or pooled samples (e.g., fecal samples) that cannot be traced or attributed to a single infected animal. Additionally, reporting of clinical signs is inconsistent and variable, making it challenging to determine the actual frequency of clinical signs associated with infection. We therefore reviewed the literature to assess reported clinical signs and virus detection during SARS-CoV-2 infection in tigers and lions [7-9, 16-28]. Additionally, we describe a SARS-CoV-2 outbreak at Denver Zoo, Colorado in late 2021, in Amur tigers (*Panthera tigris altaica*), African lions (*Panthera leo*), and spotted hyenas (*Crocuta crocuta*) (the first and only known cases in this species) [29]. We measured viral load and kinetics in animals over the course of three months, cultured infectious virus from nasal swab samples, and evaluated levels of neutralizing seroantibodies in a subset of animals. Genetic analyses reveal that all animals were infected with a SARS-CoV-2 lineage circulating at less than 1% in local human populations at that time. This study highlights the importance of the human-animal-environment interface and the potential impact of human pathogens on animals.

## Results

### SARS-CoV-2 outbreaks in United States conservatory animals

Since the beginning of the pandemic, SARS-CoV-2 has been detected in many species of conservatory animals. According to USDA Animal and Plant Health Inspection Service (APHIS) data, from April 1, 2020 to November 15, 2023, 170 conservatory animals had documented SAR-CoV-2 infections, with most occurring late ’21 to early ’22, corresponding with the wave of Delta infections (**Figure 1a, Supplemental Table 1**). The majority of these animals were nondomestic felids, with tigers and lions making up 65% of total infections, and 11 other species making up the remainder of the cases (**Figure 1b**). SARS-CoV-2 infections in conservatory animals were reported in many states, with some states reporting infections in multiple species (**Figure 1c**).

**Figure 1.**
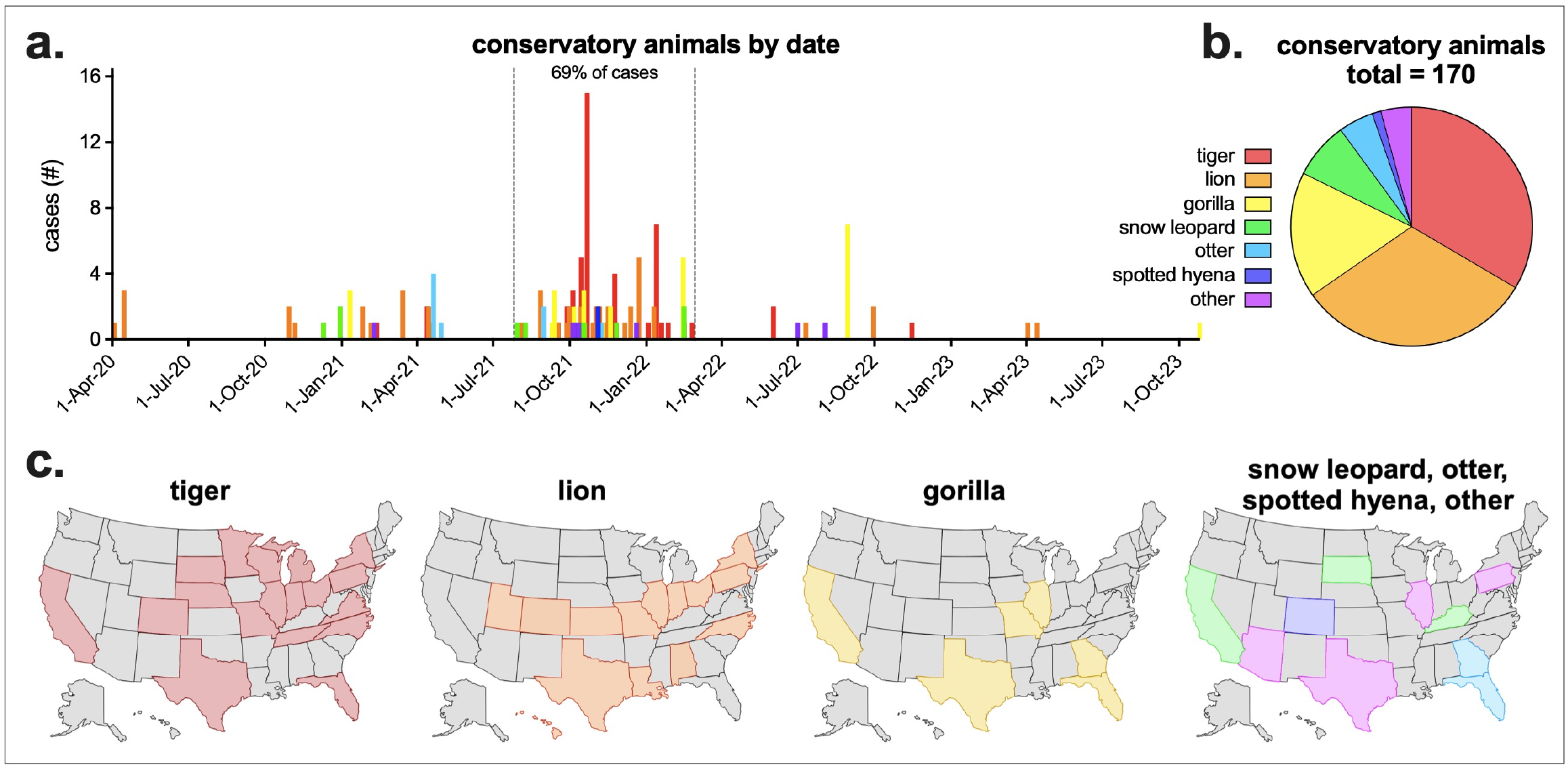
SARS-CoV-2 in United States Conservatory Animals. **a)** USDA Animal and Plant Health Inspection Service (APHIS) reported SARS-CoV-2 conservatory animal cases by confirmed date and species. Other animals include cougar, fishing cat, binturong, coati, lynx, squirrel monkey and mandrill. Sixty-nine percent of all cases occurred between July 26, 2021 and February 28, 2022. **b)** Total confirmed conservatory animal cases by animal type (n = 170). **c)** Confirmed tiger, lion, gorilla and additional animal cases by state. APHIS data downloaded November 15, 2023. Complete APHIS data provided in **Supplementary Table 1**.

### Meta-analyses of SARS-CoV-2 infections in tigers and lions

Due to the high number of reported tiger and lion infections in US conservatory animals, we performed a literature review of all published tiger and lion SARS-CoV-2 infections. There were 16 studies describing tiger/lion infection (not including retrospective serosurveys), including multiple species of both tigers and lions, with animals from multiple countries across the world (**Supplemental Table 2, Figure 2a & b**). There was a mix of male and female infections and a range of ages (2-20 years of age), with an average age of 9.6 years (**Figure 2c & d**). Not all studies reported clinical signs or virus detection (either nasal/oral, and/or fecal), but of those that did, there was a large range in the length of time clinical signs and virus detection were observed (**Figure 2e**). Of the 72 lions and tigers reported with clinical signs, there were 16 unique clinical signs, divided into four broad categories: general, nasal, respiratory, ocular (**Table 1**). Eighty-eight percent of animals exhibited respiratory clinical signs, with coughing being the most common (71% of all animals; 95% of tigers and 62% of lions). Most clinical signs were similarly detected in both tigers and lions (e.g., hyporexia/anorexia, nasal discharge, sneeze, etc.), while others were more frequently detected in one species versus the other (e.g., wheezing in 40% of tigers but only 6% of lions).

**Figure 2.**
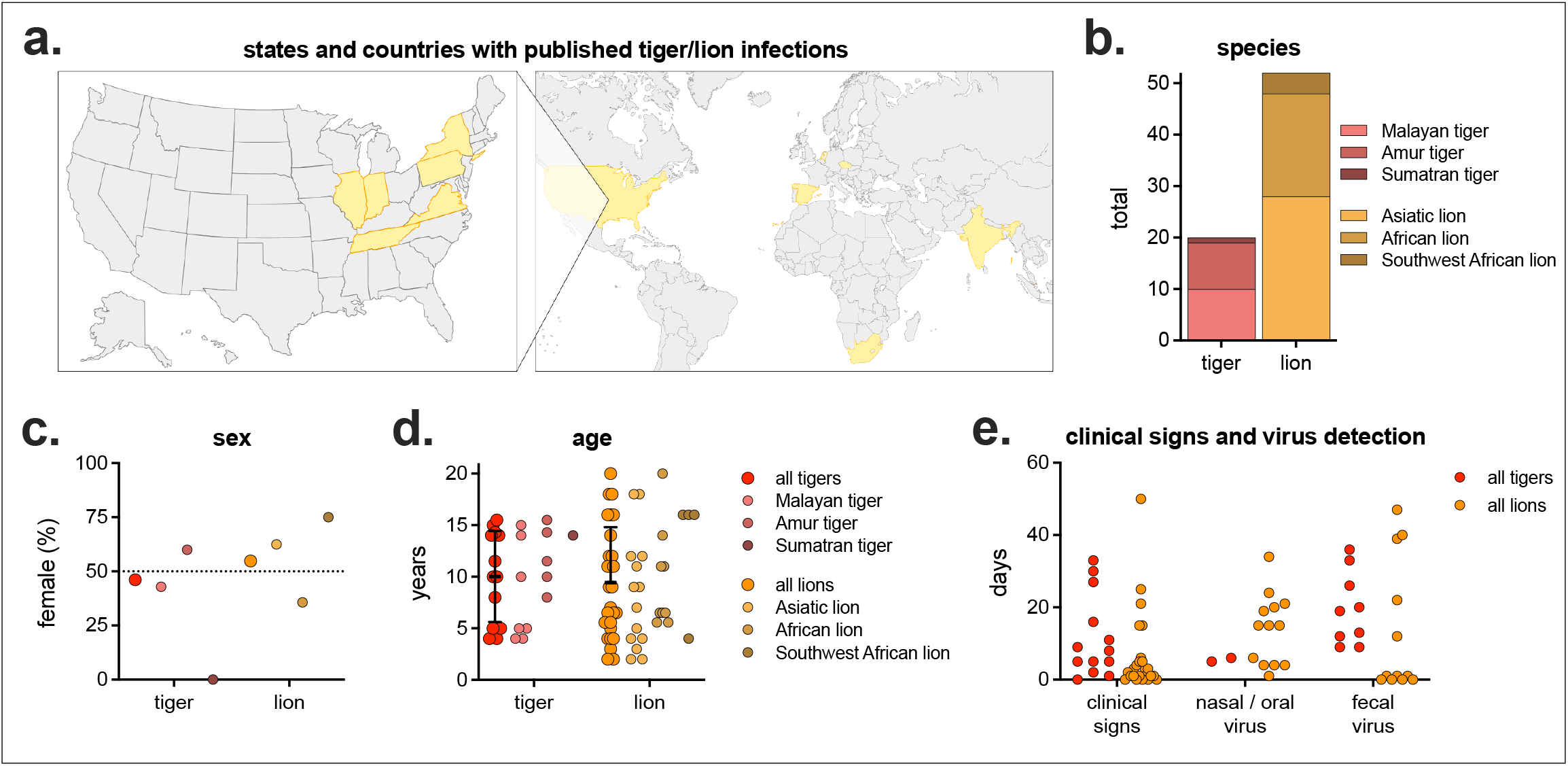
Meta-analyses of published tiger and lion SARS-CoV-2 infections. **a)** States and countries with published tiger/lion infections (n = 13 states/countries, complete list in **Supplemental Table 2**). **b)** Number and subspecies of tiger and lion. **c)** Percent female and **d)** age by species. **e**) Number of days of clinical signs, virus detection in nasal/oral samples (e.g., nasal swab, nasopharyngeal swab, saliva, etc.), and virus detection in fecal samples (feces or fecal swab). Not all studies reported sex, age, clinical signs and virus detection.

**Table 1.** Clinical sign type and frequency in published tiger and lion SARS-CoV-2 infections.

|  | all animals |  | tigers |  | lions |  |
| --- | --- | --- | --- | --- | --- | --- |
|  | % | (n/n) | % | (n/n) | % | (n/n) |
| any clinical sign | 89 | (64/72) | 95 | (19/20) | 87 | (45/52) |
| general | 43 | (31/72) | 45 | (9/20) | 42 | (22/52) |
| fever | 4 | (3/72) | 0 | (0/20) | 6 | (3/52) |
| hyporexia / anorexia | 35 | (25/72) | 45 | (9/20) | 31 | (16/52) |
| increased water intake | 4 | (3/72) | 0 | (0/20) | 6 | (3/52) |
| lethargy / exhaustion | 22 | (16/72) | 40 | (8/20) | 15 | (8/52) |
| shivering | 1 | (1/72) | 0 | (0/20) | 2 | (1/52) |
| nasal | 44 | (32/72) | 35 | (7/20) | 48 | (25/52) |
| mild epistaxis | 4 | (3/72) | 0 | (0/20) | 6 | (3/52) |
| nasal discharge | 31 | (22/72) | 15 | (3/20) | 37 | (19/52) |
| sneeze | 21 | (15/72) | 20 | (4/20) | 21 | (11/52) |
| respiratory | 88 | (63/72) | 95 | (19/20) | 85 | (44/52) |
| cough | 71 | (51/72) | 95 | (19/20) | 62 | (32/52) |
| increased respiratory rate with distress | 10 | (7/72) | 0 | (0/20) | 13 | (7/52) |
| intermittent upper respiratory sounds | 1 | (1/72) | 5 | (1/20) | 0 | (0/52) |
| labored breathing / dyspnea | 4 | (3/72) | 10 | (2/20) | 2 | (1/52) |
| respiratory symptoms | 13 | (9/72) | 0 | (0/20) | 17 | (9/52) |
| wheezing | 15 | (11/72) | 40 | (8/20) | 6 | (3/52) |
| ocular | 2 | (2/72) | 0 | (0/20) | 4 | (2/52) |
| exposed nictitans | 1 | (1/72) | 0 | (0/20) | 2 | (1/52) |
| ocular discharge | 1 | (1/72) | 0 | (0/20) | 2 | (1/52) |

Studies included 72 infected animals with reported clinical signs. Complete list of studies, and overview of animals provided in Supplemental Table 2.

### Denver Zoo SARS-CoV-2 outbreak

On October 7^th^, 2021, animal care staff at Denver Zoo observed coughing, sneezing, lethargy and nasal discharge in both 11-year-old Amur tigers (*Panthera tigris altaica*) (**Table 2**). Nasal swabs were collected and tested positive for SARS-CoV-2 viral RNA by qRT-PCR at the Colorado State University Veterinary Diagnostic Laboratory in Fort Collins. On October 14^th^, animal care staff observed the same clinical signs in both of their prides of African lions (*Panthera leo melanochaita*). Eleven lions ranging in age from 1 to 9 years all exhibited signs, and vRNA was detected in all (**Table 2**). On October 28^th^, two spotted hyenas (*Crocuta crocuta*), 22 and 23 years of age, exhibited mild clinical signs, including slight lethargy, some nasal discharge and occasional coughing, and SARS-CoV-2 vRNA was detected in both hyenas (**Table 2**). The lions and hyenas were cycled separately through the same two rotational yards in Predator Ridge; however, the two lion prides and the two hyena clans never shared a space or interacted (**Figure 3**). Both tigers were housed together in The Edge, a ∼1-acre habitat approximately 800 feet away from Predator Ridge.

**Figure 3.**
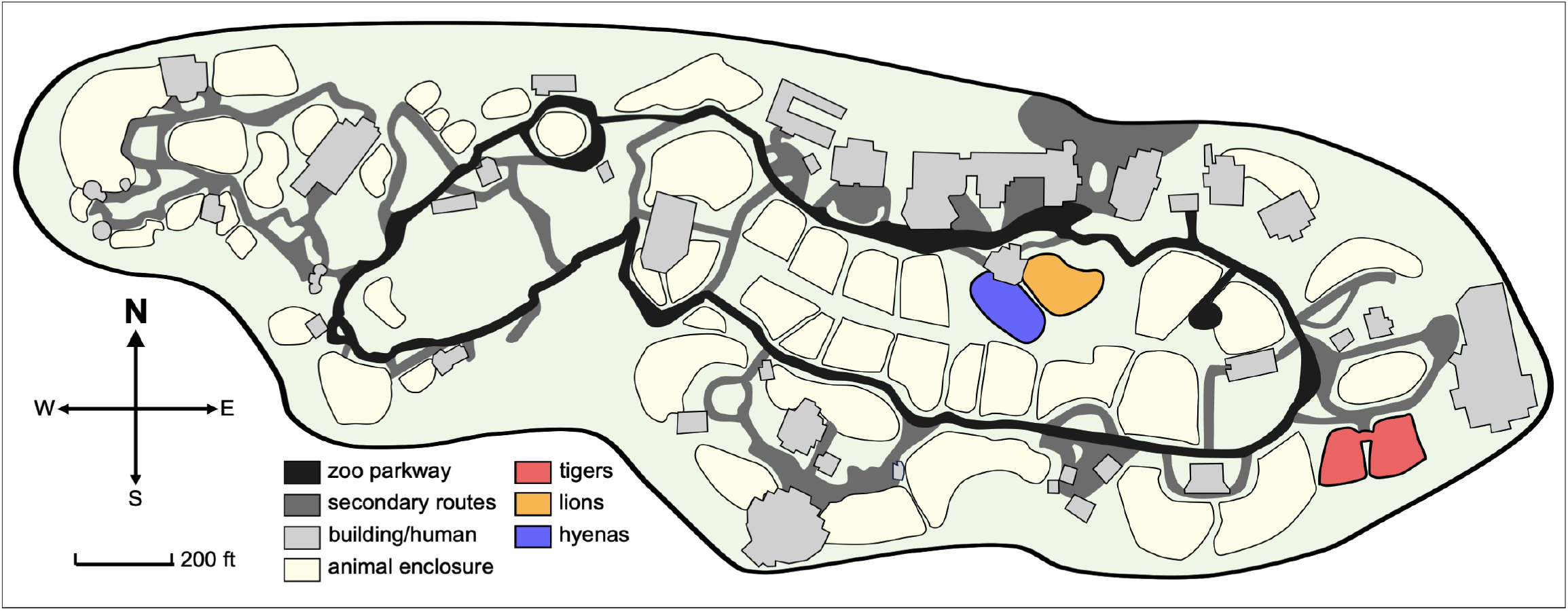
Denver Zoo map. Denver Zoo is an 80-acre zoological garden containing diverse animal habitats. Enclosures of animals that tested positive (tigers, lion and hyenas) are highlighted in red, orange and blue respectively. Tigers are housed in The Edge, an approximately 1-acre habitat designed to replicate Siberia. Lions and hyenas are cycled through two rotational yards in Predator Ridge, a nearly 5-acre space designed to replicate the landscape of the African savanna.

**Table 2.**
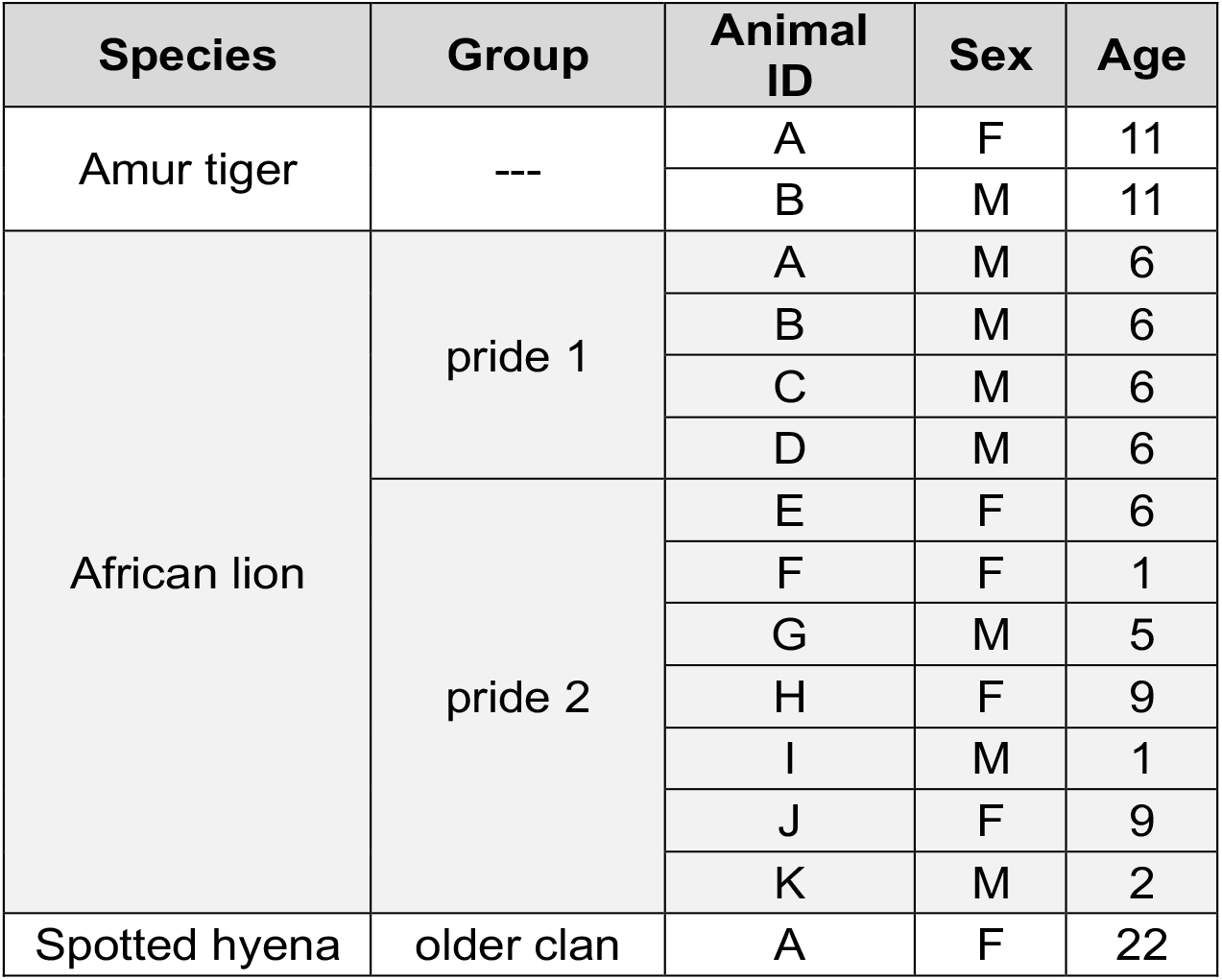
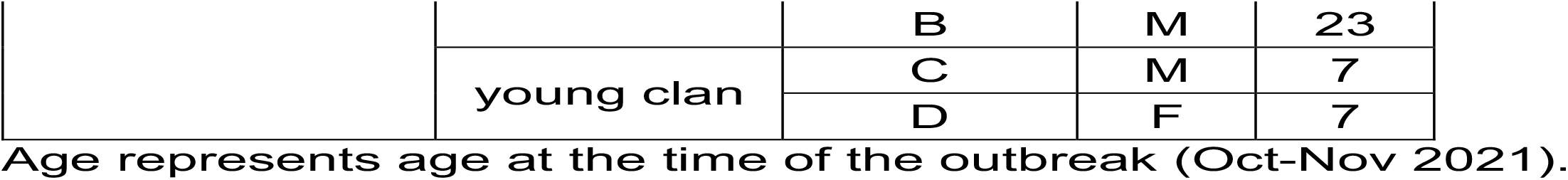
Denver Zoo animals in the study.

| Species | Group | Animal ID | Sex | Age |
| --- | --- | --- | --- | --- |
| Amur tiger | --- | A | F | 11 |
|  |  | B | M | 11 |
| African lion | pride 1 | A | M | 6 |
|  |  | B | M | 6 |
|  |  | C | M | 6 |
|  |  | D | M | 6 |
|  | pride 2 | E | F | 6 |
|  |  | F | F | 1 |
|  |  | G | M | 5 |
|  |  | H | F | 9 |
|  |  | I | M | 1 |
|  |  | J | F | 9 |
|  |  | K | M | 2 |
| Spotted hyena | older clan | A | F | 22 |
|  |  | B | M | 23 |
|  | young clan | C | M | 7 |
|  |  | D | F | 7 |

Age represents age at the time of the outbreak (Oct-Nov 2021).

### SARS-CoV-2 viral RNA in tigers, lions and hyenas

After initial positive tests, both tigers were sampled 5 more times over the course of three weeks until clinical signs resolved and three consecutive negative tests occurred (**Figure 4**). Lions and hyenas were continually sampled and tested for 3 months. Most lions appeared to resolve the infection within a month, with only negative and inconclusive (equivocal) subsequent tests and cessation of clinical signs (**Figure 4**). Almost half of the lions (five of 11), however, had recrudescent positive tests up to two months after the initial positive, and after periods of multiple consecutive negative tests (e.g., Lion B, Lion I, etc.) (**Figure 4**). Two younger hyenas (Hyenas C and D) never tested positive; however, they did have multiple inconclusive results at the same time as the older hyenas were testing positive (**Figure 4**). The first sample for all positive animals contained the highest levels of vRNA, suggesting the sample was collected at the middle to end of the course of infection (**Figure 5a**). Hyenas and tigers appeared to clear the infection/vRNA within 2-3 weeks, whereas vRNA levels decreased in all lions, then spiked again in certain individuals after multiple undetectable tests (**Figure 5a**). Lions had significantly more total vRNA compared to hyenas and tigers (**Figure 5b**).

**Figure 4.**
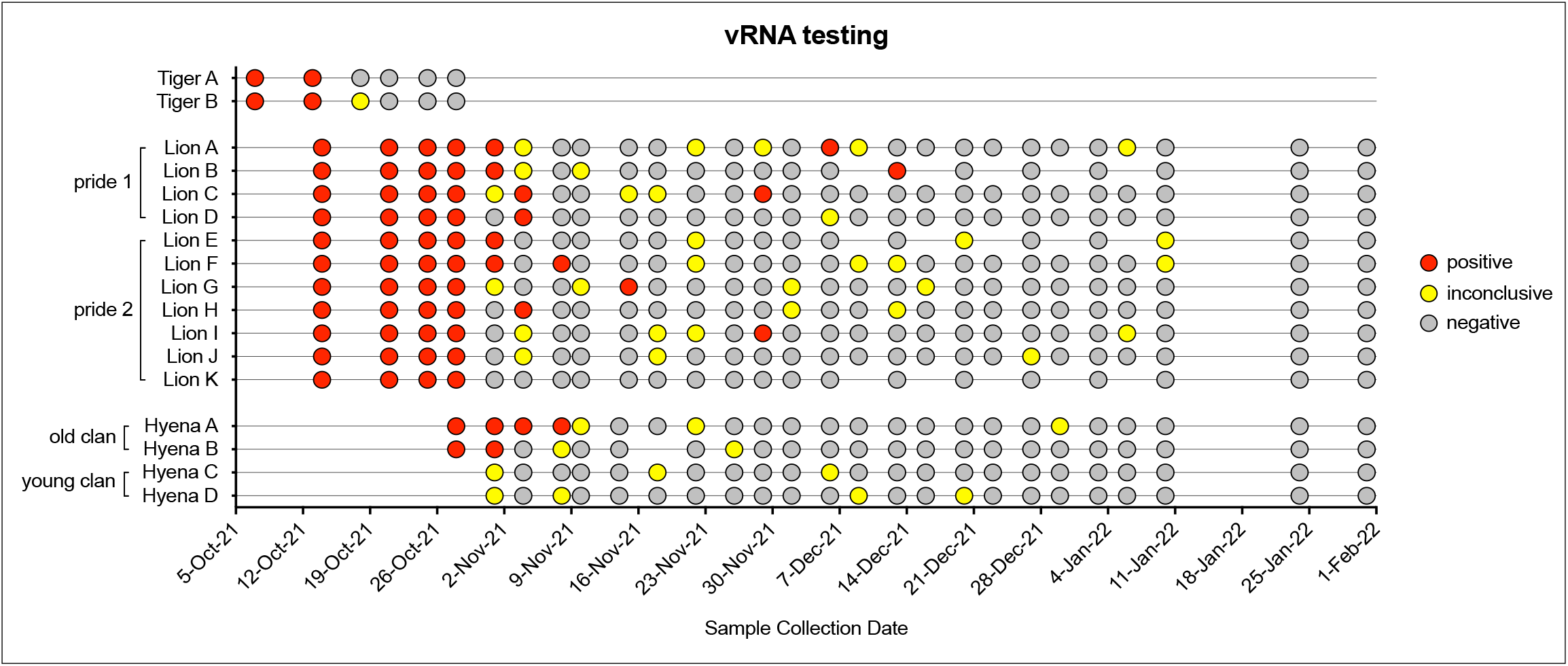
Overview of SARS-CoV-2 qRT-PCR diagnostic testing results. Voluntary nasal swabs were collected, viral RNA extracted, and tested for three SARS-CoV-2 targets (N gene, ORF1ab, and S gene) via multiplex qRT-PCR. Samples were classified as positive (red), inconclusive (yellow), or negative (gray) for SARS-CoV-2.

**Figure 5.**
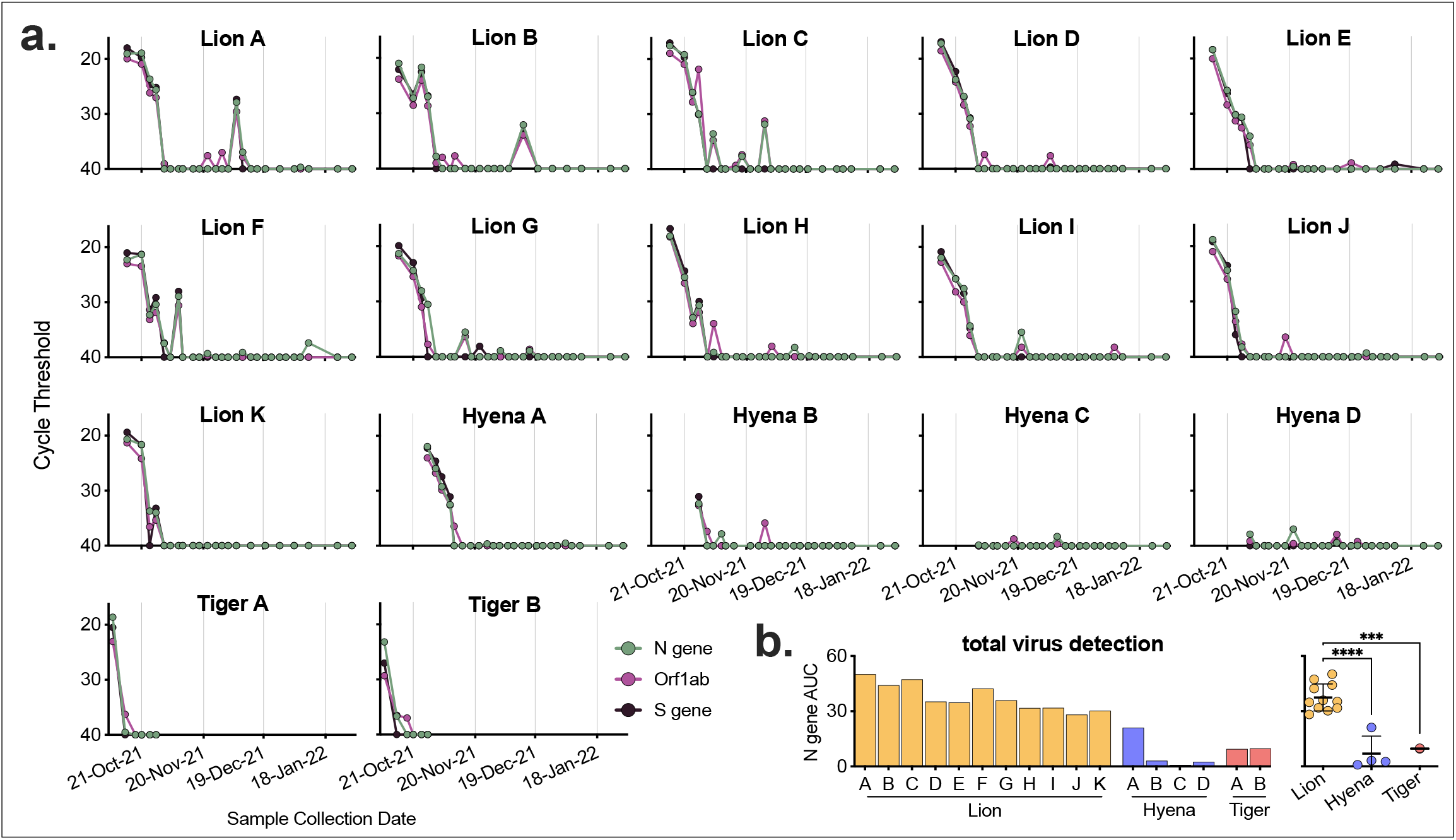
Levels of SARS-CoV-2 RNA in lion, hyena and tiger nasal swabs. **a)** Voluntary nasal swabs were collected, viral RNA extracted, and tested for three SARS-CoV-2 targets (N gene, ORF1ab, and S gene) via multiplex qRT-PCR. Undetected samples plotted at cycle threshold of 40. **b)** Total area under the curve for N gene was calculated for each animal and compared across animals. One-way ANOVA with Tukey’s multiple comparisons test (***p<0.001,****p<0.0001).

### Infectious SARS-CoV-2 in nasal swab samples

We tested a subset of nasal swab samples from lions and hyenas for infectious virus via standard plaque assay. Samples collected early in infection (Lion C and Hyena A) contained high levels of infectious virus, whereas the majority of samples tested past 2 weeks from initial positive PCR rarely contained infectious virus (**Figure 6a**). Viral titer was correlated to both timing – the closer to the first vRNA positive test, the higher the level of infectious virus – and levels of vRNA (**Figure 6b & c**).

**Figure 6.**
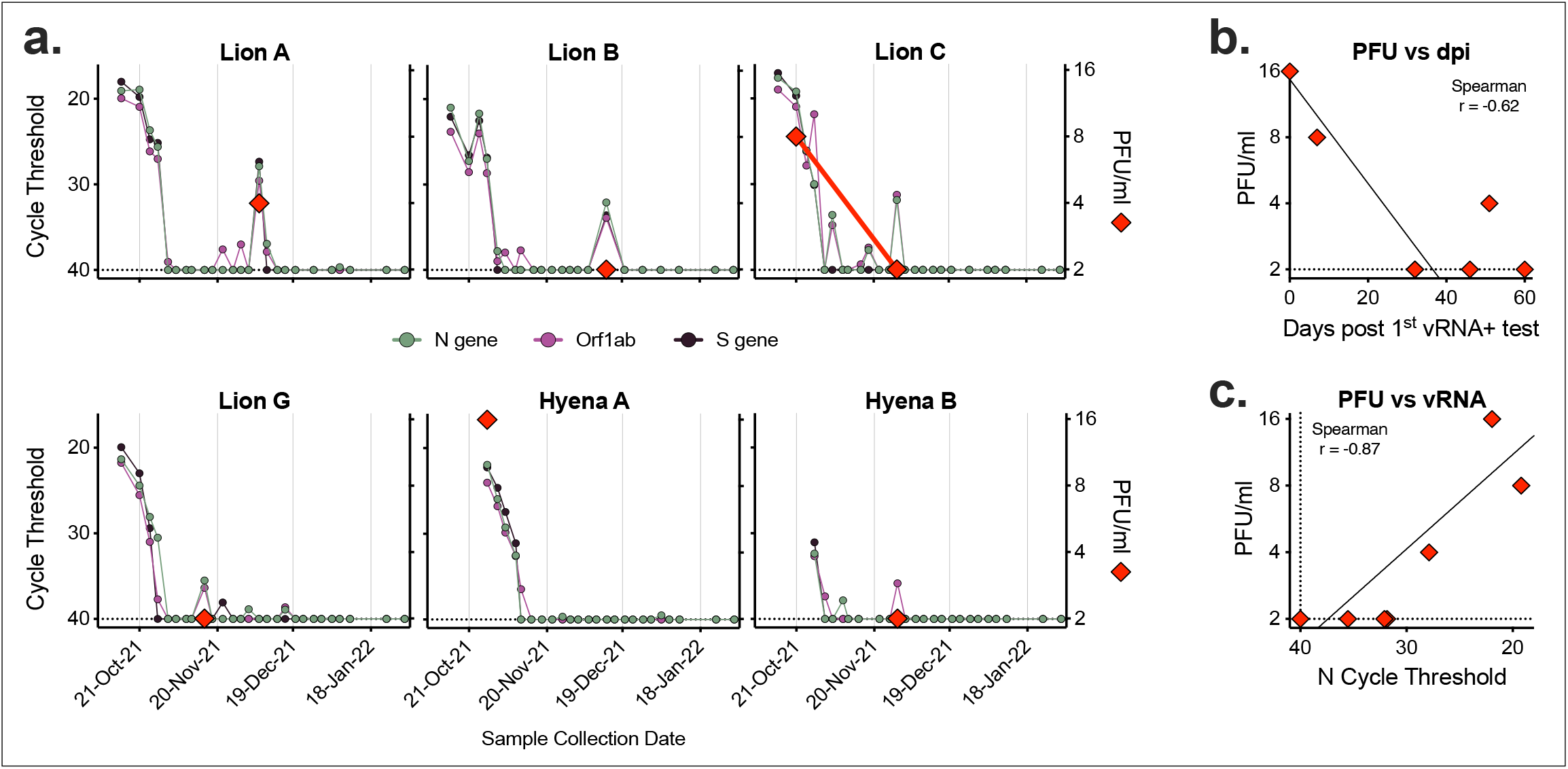
Infectious virus detected in samples with high levels of vRNA. Seven nasal swab samples from six animals were assayed for levels of infectious virus via plaque assay. **a)** Cycle threshold of three SARS-CoV-2 targets (N gene, ORF1ab and S gene, left axis) were compared to levels of infectious virus (PFU/ml, right axis). Relationship between level of infectious virus and **b)** number of days since first positive vRNA test, and **c)** N gene cycle threshold. Dashed lines represent limits of detection.

### Generation of neutralizing antibodies to SARS-CoV-2

No Denver Zoo animals in this study were vaccinated against SARS-CoV-2 prior to the outbreak. Therefore, months after animals cleared the SARS-CoV-2 infection, we tested blood from a subset of individuals for the presence and level of infection-elicited neutralizing antibodies. Both lions C and J generated high levels of neutralizing antibodies specific to SARS-CoV-2 (**Figure 7**). Despite the advanced age of Hyena B (23 years) and possibility of immunosenescence, we detected a robust neutralizing antibody response comparable to that of both lions (**Figure 7**).

**Figure 7.**
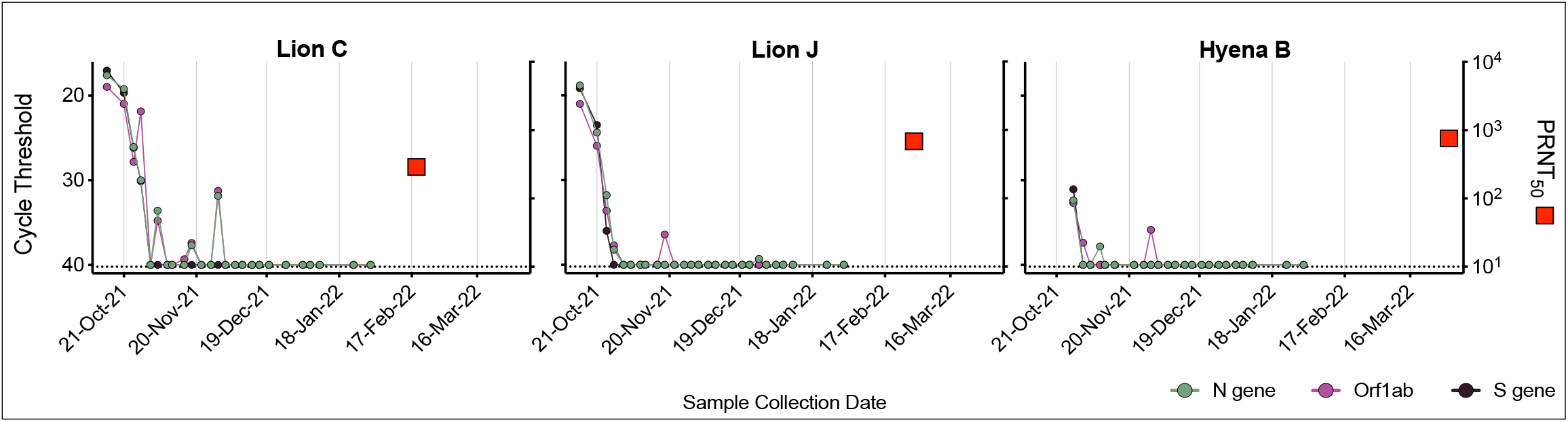
Animals generated robust neutralizing antibodies post-infection. Serum collected from three animals (Lion C, Lion J and Hyena B) approximately 4-5 months after first vRNA positive test were assayed for neutralizing antibodies using a plaque reduction neutralization test (PRNT). Cycle threshold of three SARS-CoV-2 targets (N gene, ORF1ab and S gene, left axis) were compared to PRNT_50_ (serum dilution factor required to neutralize 50% of virus, right axis). Dashed lines represent limits of detection.

### All animals were infected with a rare SARS-CoV-2 variant

SARS-CoV-2 whole genome sequencing of the first positive sample from all lions and one hyena were performed by National Veterinary Services Laboratories, USDA. Consensus whole genome analysis revealed all sequences were nearly identical, with three unique single nucleotide polymorphisms (SNPs) between the lion sequences, and three additional SNPs in the hyena sequence (**Figure 8a**). All sequences were classified as the AY.20 lineage, a sublineage within the Delta variant. The AY.20 lineage was rare in reported human SARS-CoV-2 sequences in Colorado in October 2021, averaging only 4.4 cases a day (out of an average of 647), representing ∼0.65% of all reported sequences that month (**Figure 8b & c**). We compared this to other reported tiger/lion cases to determine the frequency of variants infecting these animals in the human population in the same location at the same time. There are 109 tiger/lion SARS-CoV-2 sequences available on GISAID from 32 different states and countries from April 2, 2020 to February 14, 2024 (**Figure 8d, Supplemental Table 3**). We identified the lineage of each tiger/lion sequence and calculated the frequency and rank of that lineage in the matched human sequences in the week prior to infection in the same location (**Supplemental Table 4**). Some tigers/lions were infected with the dominant sequence circulating in humans (number 1 rank, >50% of all sequences), whereas others were infected with lineages that made up less that 10% of total human sequences (**Figure 8e**). Denver zoo samples were at the extreme end of this range, with animals being infected with a rare lineage that was circulating in humans at extremely low levels during that time (**Figure 8e**).

**Figure 8.**
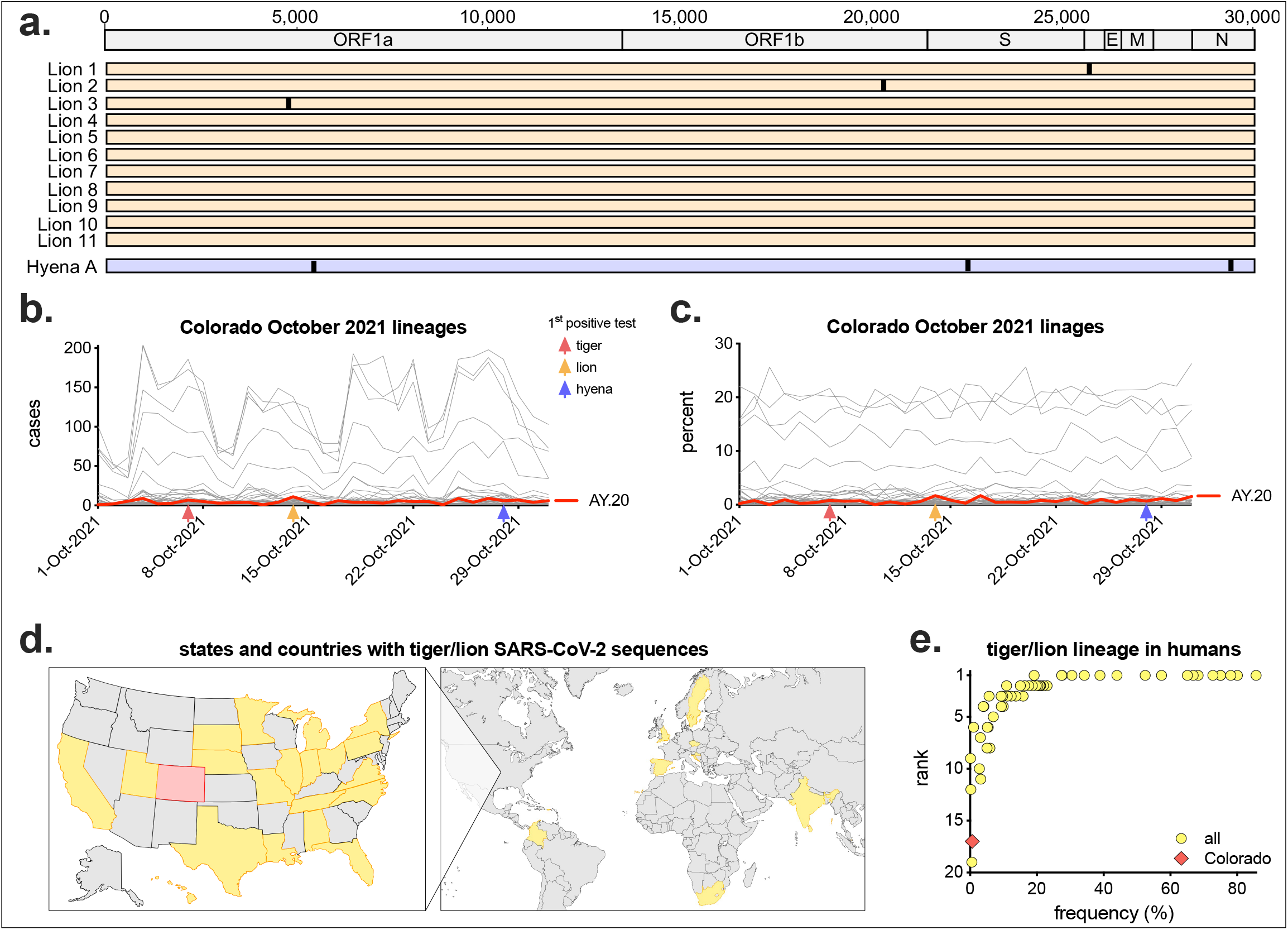
Animals were infected with AY.20, a rare variant of SARS-CoV-2. **a)** Virus from each of the 11 lions collected on October 14, 2021 and Hyena A collected on October 28, 2021 (first positive test for each animal) were sequenced and determined to be Delta lineage AY.20. Consensus whole genome sequences were aligned (numbers represent nucleotide position), with vertical lines representing single nucleotide polymorphisms between viruses. For each lineage, the **b)** number of cases and **c)** frequency as a percentage of all human SARS-CoV-2 sequences reported in Colorado in October 2021 (n = 20,051) were plotted by date. Red line shows lineage AY.20. Red, orange and blue arrows represent the date of first positive test in tigers, lions and hyenas respectively. **d)** Map of states and countries with full genome sequences from tigers and lions (n=109, Colorado in red). **e)** For each tiger/lion sequence, the frequency and rank of that lineage in humans in the same state/country in the week prior to infection was calculated (complete list in **Supplemental Table 4**).

## Discussion

It is well established that many animal species, especially cats, are susceptible hosts for SARS-CoV-2 [12]. Natural infections and experimental studies in domestic cats, and natural infections in big cats in zoological institutions have revealed a broad range in infection kinetics, clearance, and common clinical signs. In this study, we provide a detailed overview of a SARS-CoV-2 outbreak that occurred at Denver Zoo beginning in October 2021, affecting all tigers, lions and hyenas in residence, none of which were vaccinated against SARS-CoV-2 at the time of infection due to inability to obtain the vaccine until after the outbreak. While our results in tigers and lions were consistent with other published studies, our repeated sampling of individuals over months provided substantially higher resolution of viral replication and clearance kinetics and demonstrated clear viral recrudescence after many weeks of consecutive negative tests, which had not previously been documented.

Additionally, detection of infectious virus and seroconversion (generation of neutralizing antibodies), and sequence analyses further refine our understanding of this outbreak and infections.

qRT-PCR is a highly sensitive test and detects vRNA, including fragments of inactive virus. Particularly with an inconclusive result, detection of vRNA does not solely indicate active infection in an animal. Detection of vRNA after initial clearance has been documented in humans after natural infection, both with and without the use of oral antivirals (e.g., Paxlovid or remdesivir) [30, 31]. After weeks of undetectable levels, we detected viral RNA in some lions, and in Lion A, moderate levels of infectious virus as well. Whether this was viral rebound within each animal, or a new re-infection that spread amongst some animals is unknown; however, subsequent sequence analyses suggest the former [32]. Infections and clinical signs were mild in Denver Zoo animals and all made full recoveries. Conversely in other instances in Sweden, Ohio, and Hawaii (especially involving older animals or those with comorbidities), tigers and lions died or were euthanized due to deteriorating health and medical complications caused by SARS-CoV-2 infection [33, 34]. A recent meta-analysis of SARS-CoV-2 infections in tigers in North American zoos found that most infections were mild, but vaccinated animals were still susceptible to infection [35]. Expanded access to vaccines and treatments (e.g., antivirals, NSAIDs, fluids, etc.) can continue to reduce morbidity and mortality in conservatory animals.

Our study is the first and only documented case of SARS-CoV-2 infections in hyenas [29]. This reveals that despite extensive viral surveillance and experimental infections in animals, a full list of potential SARS-CoV-2 hosts remains unknown. Hyenas group phylogenetically with Feloidea, the ‘cat branch’ of the Carnivora order, making them more similar to Felidae (cats, lions, tigers, lynxes, etc.), Viverridae (civets, binturong, etc.), and Herpestidae (mongooses, meerkats, etc.), than Canidae (dogs, foxes, wolves, etc.) [36]. Many of these Feloidea species (e.g., cats, lions, tigers, binturong, civets, fishing cats, etc.) are susceptible to SARS-CoV-2 and related coronaviruses (SARS-CoV, Civet SARS-CoV, etc.) [16, 37, 38], suggesting other unstudied Feloidae species may be susceptible, and potentially serve as natural reservoirs as well [39, 40]. Despite the important One Health implications, any potential benefit of detecting SARS-CoV-2 infections in wild animals (especially novel hosts), and subsequent impact on risk assessment and management, must be balanced with the challenges and cost of such surveillance efforts [41].

Comparison of viral sequences from the first positive test from all lions and one hyena revealed almost identical viruses, demonstrating they were all infected with the same variant. Importantly, this variant was present in humans in Colorado at this time at incredibly low levels (∼0.65% of all sequences). These data suggest there was a single spillover event from an infected individual, into animals (likely tigers as they were the first to show clinical signs), then spread within and across species. While the lions and hyenas rotate through shared spaces, and transmission between them is thus feasible, it remains unknown how virus from tigers was transmitted to lions/hyenas, as their enclosures are over 800 feet apart, with no shared staff working at both locations. It is possible that an intermediate host present at Denver Zoo, such as squirrels or mice [42], transmitted the virus between locations, or that personnel or equipment served as a fomite. Additionally, while it could be possible a highly infectious zoo visitor with this variant independently transmitted the virus to both tigers and lions, in-depth sequence analyses by Bashor *et al*. demonstrate that virus present in all lions and hyenas contains mutations present in the tiger sequences, refuting this hypothesis [32]. These analyses also reveal novel shared mutations and unique rapid evolution within and across species over the course of infection [32].

There remain many unknowns regarding SARS-CoV-2 infections in wild and conservatory animals, and the potential role they play as viral reservoirs, sources of novel variants, and in spillback transmission of virus back into human populations. Pathogen surveillance of wild and conservatory animals requires extensive time, cost and resources [41]. Modeling and *in vitro* assays testing compatibility of viral proteins and host receptors have revealed potentially susceptible species that have not yet been documented in nature, including alpacas and pandas [43, 44]. Expanded molecular and modeling assays, in combination with increased targeted surveillance of vulnerable animals (e.g., conservatory animals with comorbidities) and those in close contact with humans (e.g., farmed mink) can improve our ability to predict and detect infections. Additionally, a collaborative One Health approach can further protect the environment, animals and humans.

## Materials and Methods

### Animals

Denver Zoo exhibits Amur (Siberian) tigers (*Panthera tigris altaica*), African lions (*Panthera leo melanochaita*) and spotted hyenas (*Crocuta crocuta*) (**Table 2**). Some of the animals had pre-existing comorbidities prior to the SARS-CoV-2 outbreak. The four lions in pride 1 were brothers (all 6 years old), and all had medical histories of gastrointestinal disease, but were clinically stable and not under treatment during the SARS-CoV-2 outbreak. Lion A additionally had lumbosacral disease. In pride 2, Lion G (male, 5 years old) had inflammatory bowel disease, and Lion H (female, 9 years of age) had cerebral atrophy and osteoarthritis. At the time of testing, Hyena B (male, 23 years of age) had elevated liver enzymes and a gastric foreign body removal.

### Animal care specialists

Prior to and during the time of the outbreak, individual animal care specialists cared for each of the three species, with shared specialists between lions and hyenas, but no overlap between them and tigers. Animal care specialists wore personal protective equipment when caring for animals, including gloves, coveralls, KN95 masks and face shields. Any staff that tested positive for SARS-CoV-2 quarantined and did not go to work at the zoo.

### Sample collection

Voluntary nasal swabs were obtained from all three species by trained animal care specialists. Nasal swab samples were collected by gently rubbing on the inside of the nose, then placed into a tube containing viral transport media. Voluntary blood was collected in serum collection tubes, allowed to clot at room temperature for at least 30 minutes, then centrifuged to separate into serum. Serum was heat-inactivated at 56°C for 30 minutes, then stored at 4°C.

### Viral RNA extraction and qRT-PCR

Swabs were transported to the Colorado State University Veterinary Diagnostic Laboratory in Fort Collins, Colorado. Viral RNA was extracted using a KingFisher Flex Magnetic Particle Processer and tested for SARS-CoV-2 by quantitative reverse transcriptase PCR (qRT-PCR) using the Thermo Fisher Scientific TaqPath COVID-19 combo kit, targeting three viral targets (ORF1ab, N gene and S gene), and an MS2 positive control [45]. qRT-PCR was performed on an Applied Biosystems 7500 Fast qPCR instrument and analyzed using Applied Biosystems COVID-19 Interpretive Software and individual review of each amplification plot. The threshold was set at 10% of the plateau of the positive amplification control. Results were classified as positive, negative, or inconclusive using the below criteria (if MS2 was not detected, result is invalid, and assay repeated). Because the S gene target was not detectable for some SARS-CoV-2 variants, detection of the S gene was not required to determine a positive result.

| ORF1ab | N gene | S gene | Result |
| --- | --- | --- | --- |
| NEG (no exponential amplification) | NEG (no exponential amplification) | NEG (no exponential amplification) | negative |
| ORF1ab and N targets +/- S target positive with one or more target with Ct < 38 |  |  | positive |
| Detectable exponential amplification in one, two or three targets but does not meet criteria to be determined positive |  |  | Inconclusive (equivocal) |

### Virus plaque assay

Vero cells (ATCC-81) were maintained in Dulbecco modified Eagle medium (DMEM) with 10% fetal bovine serum (FBS) and 10 units/mL penicillin, 10 μg/mL streptomycin, and 2.5 μg/mL amphotericin B at 37°C and 5% CO_2_. Cells were plated one day prior to infection in 12-well tissue culture plates. Nasal swab samples were serially diluted in infection media (growth media with 1% FBS), then inoculated onto cells for 1 hour at 37°C. After incubation, cells were overlaid with 0.6% tragacanth medium, incubated for 2 days at 37°C, and fixed and stained with 30% ethanol and 0.1% crystal violet. Plaques were counted manually.

### Plaque reduction neutralization assay

A standard plaque reduction neutralization test (PRNT) was performed using SARS-CoV-2 virus (2019-nCoV/USA-WA1/2020 strain) [46]. Heat-inactivated, diluted serum samples were mixed with virus, incubated for 1 hour at 37°C, added to Vero cells, and incubated for an additional hour at 37°C. Cells were then overlaid with tragacanth medium and incubated for 2 days at 37°C. Cells were fixed and stained with ethanol and crystal violet, and plaques counted manually.

### Lion and hyena SARS-CoV-2 sequence analysis

Denver Zoo lion and hyena samples were sequenced by the USDA Animal and Plant Health Inspection Service (APHIS) using an Illumina MiSeq and IRMA v0.6.7; DNAStar SeqMan NGen v14.1.0 assembly methods. Sequences are available under GISAID accession numbers EPI_ISL_6088015, EPI_ISL_6088016, EPI_ISL_6088018, EPI_ISL_6088025, EPI_ISL_6088026, EPI_ISL_6088034, EPI_ISL_6088040, EPI_ISL_6088041, EPI_ISL_6088044, EPI_ISL_6088046, EPI_ISL_6088051, and EPI_ISL_6810900. The lion sequences are deidentified and unable to be matched to Denver Zoo animal IDs.

### SARS-CoV-2 variant analysis

For Colorado analyses, all complete sequences, low coverage excluded, from humans from October 1^st^ to the 31^st^, 2021 in the state of Colorado were downloaded from GISAID (n = 20,051). Total counts of each lineage, and percent of each lineage by day were determined. For tiger and lion analyses, all complete sequences, low coverage excluded, from all Panthera were downloaded from GISAID (February 14, 2024, n = 118). Of those, 109 sequences came from tigers and lions (9 snow leopard sequences were excluded from analyses). For each sequence, matching human sequences (complete, low coverage excluded), from the same region were downloaded for the week prior to tiger/lion infection (**Supplemental Tables 3 and 4**). For each tiger/lion sequence lineage, the frequency and rank of the same lineage in the human sequences were determined.

### Statistical analyses

Statistical analyses were performed using GraphPad Prism Version 10.2.3. Statistical tests and details are provided in figure legends.

## Acknowledgements

ENG was supported by funding to Verena (viralemergence.org) from the U.S. National Science Foundation, including NSF BII 2021909 and NSF BII 2213854. LB was supported by the National Institute of Allergy and Infectious Diseases of the National Institutes of Health under Award Number T32AI162691. The research reported in this publication was supported by the Colorado State University Veterinary Diagnostic Laboratories, Colorado State University College of Veterinary Medicine and Biomedical Sciences Research Council Award and Colorado State University’s Office of the Vice President for Research’s ‘Accelerating Innovations in Pandemic Disease’ initiative, made possible through support from The Anschutz Foundation. We gratefully acknowledge the many collaborators on and contributors to this project, especially all the dedicated staff at Denver Zoo, including the animal care specialists. We thank the staff of the CSU Veterinary Diagnostic Laboratories who performed the diagnostic testing. We also thank Ellie Graeden for discussion and suggestion on the results of the study.

## Author contributions

ENG – data curation, formal analysis, investigation, visualization, writing-original draft, writing-review & editing; LB – formal analysis, writing-original draft, writing-review & editing; KE – data curation, formal analysis, investigation, resources; LC, KS and JL – investigation, resources; SVW – supervision, writing-review & editing; JGJ – conceptualization, resources, writing-review & editing; KP – conceptualization, funding acquisition, project administration, resources, supervision, writing-review & editing; GDE – conceptualization, supervision, writing-review & editing.

## Figure legends

**Supplemental Table 1. APHIS conservatory animals**. Data downloaded November 15, 2023.

**Supplemental Table 2. Overview of published studies of tiger and lion SARS-CoV-2 infections**.

**Supplemental Table 3. States and countries with tiger and lion SARS-CoV-2 sequences**.

**Supplemental Table 4. Tiger and lion lineage frequency and rank in matched human sequences**.

